# Atypical flagella assembly and haploid genome coiling during male gamete formation in *Plasmodium*

**DOI:** 10.1101/2023.05.17.540968

**Authors:** Molly Hair, Flavia Moreira-Leite, David Ferguson, Mohammed Zeeshan, Rita Tewari, Sue Vaughan

## Abstract

*Plasmodium* spp. sexual reproduction occurs within the *Anopheles* mosquito and is essential for gametogenesis and onwards transmission to mammalian hosts. Upon activation, the male *P. berghei* gametocyte undergoes three rounds of inter-nuclear mitosis and assembles eight basal bodies and axonemes around the nucleus prior to ex-flagellation, resulting in 8 flagellated male gametes in 12-15 minutes. However there is little understanding of the 3D organisation of this rapid process of male gametogenesis. In this study we used serial block face scanning electron microscopy (SBF-SEM) and cellular electron tomography (ssET) of *P. berghei* microgametocytes to examine the 3D architecture of key structures during male gamete formation. Our data has revealed an exquisite organisation of axonemes coiling around the nucleus in opposite directions forming a central ‘axoneme band’ in microgametocytes. Furthermore, we discovered that the nucleus of microgametes is tightly coiled around the axoneme in an exquisitely complex structure whose formation starts before microgamete emergence during ex-flagellation. Our discoveries of the detailed 3D organisation of the flagellated microgamete and the haploid genome highlights some of the atypical mechanisms of axoneme assembly and haploid genome organisation during male gamete formation in the malaria parasite *Plasmodium spp*.

## Introduction

The prerequisite for the transmission of *Plasmodium* spp. from an infected mammalian host to *Anopheles* mosquito vector relies upon the parasite switching from asexual to sexual reproduction. This maturation occurs when an asexual schizont differentiates into a sexually committed merozoite which gives rise to immature male and female within the blood of the intermediate host gametocytes (Josling & Llinás, 2015; R. E. Sinden, 1983; Venugopal et al., 2020). Only on ingestion of blood meal by an *Anopheles* mosquito do these gametocytes undergo development to produce both male (microgamete) and female (macrogamete) (Josling & Llinás, 2015; Kuehn & Pradel, 2010; R. Sinden et al., 2010). Within the mosquito’s midgut, environmental changes of pH, the presence of xanthurenic acid and a drop in temperature leads to stimuli activating the micro-and macrogametocytes. Post activation microgametocytes undergo three rounds of atypical intra-nuclear mitosis from haploid (1N) to octaploid (8N) and assemble eight axonemes within the cytoplasm of the microgametocyte without intraflagellar transport. The microgametocyte then undergoes ex-flagellation to release eight haploid motile microgametes capable of fertilising macrogametes with 15 minutes (R. Sinden et al., 1978, 2010; R. E. Sinden, 1982).

Coordinating this event are bipartite microtubule organising centres (MTOCs) each composed of an acentriolar MTOC (called a nuclear pole NP), which lies on inner nucleoplasmic side of the nuclear envelope and is responsible for spindle microtubule formation and a centriolar MTOC (called a basal body BB) associated with an axoneme on the outer side of the nuclear envelope. The cellular ultrastructure of nuclear poles (NPs) and basal bodies (BBs) have previously been observed by electron microscopy as two separate electron-dense structures, bridging between the nuclear and cytoplasmic sides of the nuclear envelope (R. Sinden et al., 2010; R. E. Sinden, 1982; Zeeshan et al., 2022). During the early stages of microgametocyte activation (1-3 minutes), the inner and outer MTOCs work initially to coordinate mitotic division and to nucleate eight axonemes with rapid axoneme assembly occurring within the cytoplasmic region around the single nucleus (Guttery et al., 2022; R. Sinden et al., 1978; R. E. Sinden, 1983; Zeeshan et al., 2022). By 6-8 mins post-activation most axonemes have formed and by 8-10 mins ex-flagellation has begun with basal bodies protruding first out of the cell membrane each one bringing with it an assembled axoneme and a haploid genomic nucleus, the essential components of motile microgametes (Rashpa & Brochet, 2022; Zeeshan et al., 2022). The whole process is usually complete by 12-15 min (Guttery et al., 2022; R. Sinden et al., 2010; R. E. Sinden, 1983). Visualising the process of microgamete assembly and ex-flagellation by thin section transmission electron microscopy (TEM) only provides 2D views of the ultrastructure, but it is clear from live cell studies that microgamete assembly is a complex three-dimensional process (Guttery et al., 2022; R. Sinden et al., 2010). Here, we have used serial block face scanning electron microscopy (SBF-SEM) to reconstruct whole individual microgametocytes and key elements in individual microgametes. In addition, we performed serial electron microscopy tomography of individual microgametes, which revealed interesting distinctiveness of their ultrastructure. The combined volume electron microscopy data presented here improves significantly our structural understanding of the dynamic process of *Plasmodium berghei* male axoneme assembly, flagellar formation and haploid genome organisation within the microgamete.

## Results & Discussion

### Analysis of basal bodies and nuclear poles reveals heterogeneity in association during male gametogenesis

*Plasmodium berghei* purified gametocytes cells isolated from infected mice blood were activated for 15 minutes and then fixed, embedded, and imaged by SBF-SEM. This process resulted in two datasets both containing 197 individual slices (Z-slice = 100nm) (Movie 1). In order to specifically visualise the three-dimensional organisation of axonemes, microgametocytes containing assembling axonemes were chosen for further analysis. From these datasets 15 individual whole microgametocytes with their axonemes internally coiled were identified and analysed. The plasma membrane, nucleus membrane, nuclear pole MTOCs (NPs), basal body MTOCs (BBs) and axonemes were segmented for each whole cell using IMOD (Kremer et al., 1996) to produce 3D models.

*Plasmodium* basal bodies do not exhibit a defined 9 triplet microtubule ultrastructure by TEM as seen in many other organisms. Their BB structure is defined as an electron-dense mass which, on rare occasions, nine single A-tubules can be resolved on the cytoplasmic side of the nuclear envelope with the NP closely associated on the inner surface of the nuclear envelope with radiating microtubules of the spindle (Rashpa & Brochet, 2022; R.E. Sinden et al., 1976; Zeeshan et al., 2022). By following along the length of each axoneme in our 3D reconstructions, it was possible to locate the BB and NP closely associated with the envelope of the nucleus. The BB and NP are physically cross-linked through the nuclear envelope (Francia et al., 2015; R.E. Sinden et al., 1976; Zeeshan et al., 2022) and in our SBF-SEM data slices electron-dense filamentous material can be observed in most cells between the two MTOCs (Fig 1A; arrows).

**Figure 1.**
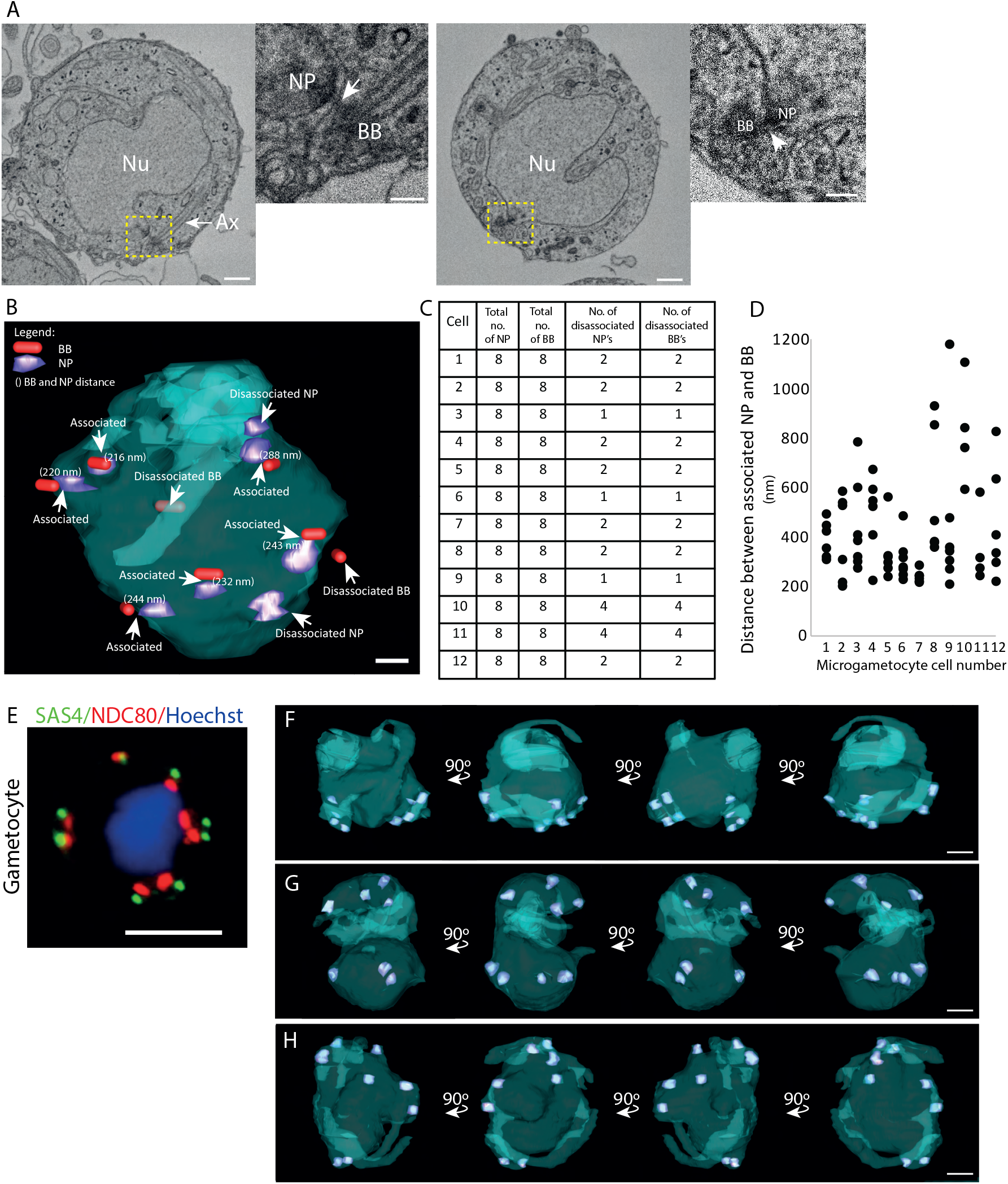
Analysis of basal bodies and nuclear poles reveals heterogeneity in association. A total of 15 microgametocyte cells 15 mins post activation were analysed and 3D reconstructed from SBF-SEM data of gametocyte populations to examine the association between basal bodies (BBs) and nuclear poles (NPs). (A) SBF-SEM data slices from two microgametocytes showing the typical ‘bipartite’ MTOC composed of a basal body (BB) and a nuclear pole (NP) linked by filamentous material (arrow). Nu, nucleus; Ax, axoneme. Scale bar = 1 μm in lower magnification images, or 200 nm in insets. (B) A 3D segmented model of a microgametocyte nucleus (blue) and ‘associated’ and ‘disassociated’ BBs (red) and NPs (purple). The distance between associated BBs and NPs were measured (nm). Scale bar = 500 nm. (C) Table showing the total number of NPs and BBs per microgametocyte cell, and the total number NPs and BBs that are disassociated (i.e. not linked forming a bipartite MTOC). (D) Dot plot graph showing the distances (nm) between the BB and the NP in associated pairs, in each microgametocyte cell. (E) Fluorescence microscopy image showing the localisation of the BB marker SAS4-GFP (green) in relation to the NP (kinetochore) marker NDC80-m-Cherry (red) in male gametocytes 8 min post-activation. Scale bar = 5 μm. (F-H) Segmented nuclei (blue) at 90-degree rotations showing the three different configurations of nuclear pole (purple) positions observed across the 15 cells. F) Nucleus showing nuclear pole pairs positioned at one side of the nucleus, (G) Nucleus showing nuclear pole pairs positioned in each ‘corner’ of the nucleus and (H) Nucleus showing nuclear pole pairs grouped together with a single pair on the opposite side of the nucleus. Scale bar = 5 μm.

Of the 15 microgametocytes analysed, 12 of these contained 8 BBs with axonemes and 8 NPs. The resolution in the remaining 3 microgametocytes was limited, therefore we were unable to clearly distinguish the NPs, but they did contain 8 BBs with axonemes. Moving through the SBF-SEM data slices of the remaining 12 microgametes where all NPs were visible allowed us to identify which BB was in close proximity to which NP. This analysis revealed that the majority (75%) of BBs and NPs were clearly associated (Fig 1B). There were BBs with assembled axonemes that were not associated with a NP and there were NPs that were not associated with a BB (Fig 1B-C). For example microgametocyte cell 1 shown in Fig 1B clearly shows 2 BBs without an NP in close proximity and 2 NPs without a BB in close proximity (Fig 1C). We measured the distance between each associated BB and NP in each of the 12 microgametocytes using the mid-point of each electron dense structure to quantify their relationship. Our data revealed the average distance between the two was 431nm (Fig 1D; n = 71). It is important for BBs and NPs to remain connected so that an axoneme can be associated with a single haploid microgamete nucleus upon ex-flagellation, but they must become disassociated at some stage so that ex-flagellation can occur. These disassociated BBs and NPs observed at this stage represent aberrant forms and are unlikely to support genome segregation into a budding microgamete. This is in agreement with data showing that male gametogenesis does not always develop correctly (Francia et al., 2015; R. Sinden et al., 2010; Straschil et al., 2010).

Next, we analysed the positioning of the 8 NPs within the nuclear envelope. As the cell undergoes three rapid rounds of DNA replication, spindle microtubules nucleate from each NP and chromosome segregate as observed with kinetochore marker NDC80 (Zeeshan et al., 2020). The basal body marker SAS4-GFP is found associated with the kinetochore marker NDC80-Cherry even at 8 minutes after activation when nuclear content is octoploid as observed by live cell imaging (Fig 1E) (Zeeshan et al., 2022). In early stages, NPs are often located in closely associated pairs. As gametogenesis progresses NP pairs are located around the circumference of the nucleus before each pair finally separate and 8 single NPs are separated around the circumference of the nuclear envelope (Rashpa & Brochet, 2022; R. Sinden et al., 2010). By SBF-SEM we found heterogeneity in the positioning of the NPs between microgametocytes at pre-exflagellation stages, which strongly suggests that NP movement is occurring whilst axonemes are forming (Fig 1F-H). In 4 cells the nuclear pole pairs were grouped on one side of the nucleus (Fig 1F). In 5 cells we saw the pairs of NPs positioned in each edge of the nucleus (Fig 1G) and in 3 cells, most of the NP pairs were grouped together with a single pair on the opposite side of the nucleus (Fig 1H). Taken together, these results show the 3D organisation of BBs and NPs and demonstrate the close association of the BB and NP during male gametogenesis.

### Axonemes coil around the nucleus in opposing directions creating an axonemal band

A hallmark stage post activation of gametocytes is the assembly of axonemes which are internally coiled around the nucleus (Fig 2A) and can clearly be seen at ∼3-4 mins after activation using live cell imaging microscopy and basal body marker SAS-GFP with an axoneme marker Kinesin8B-mcherry (Zeeshan et al., 2019). Our SBF-SEM dataset has allowed further detailed analysis of this coiling which we term an axonemal band. In the 15 microgametes analysed each had 8 axonemes internally coiled around the nucleus and in close association with one another (Fig 2B). Using the BB as an orientation marker, we discovered that axonemes do not all coil around the nucleus in the same direction. Each axoneme was segmented in yellow or blue depending on their orientation around the nucleus (Fig 2B, Movie 2). We found that the two axonemes of each BB/NP pair coil in the same direction, suggesting a level of positional constraint (Fig 2C-D). We found only one occasion where the two axonemes of a BB/NP pair were coiled in opposite directions. Taken together, these results suggest there is a level of asymmetry/chirality in the ultrastructural organisation of BB/NP/axoneme pairs in the gametocyte. This asymmetry may be determined by the BB structure or mitotic spindle organisation, ensuring that each pair coils around the nucleus rotates in a defined direction. Presumably, this could be advantageous in preventing intertwining during axonemal beating.

**Figure 2.**
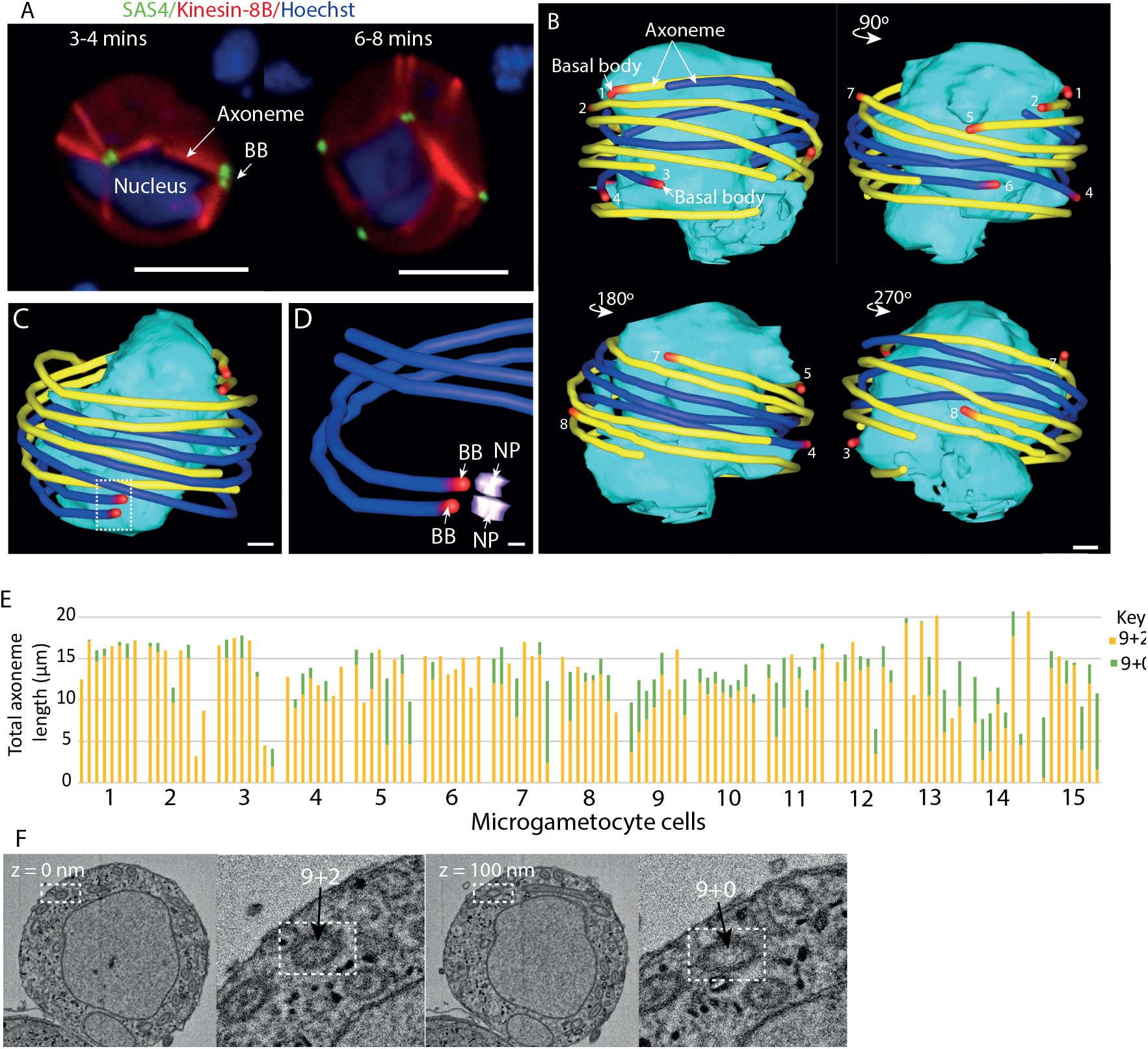
Axonemes coil around the nucleus of the microgametocyte in opposite directions creating an axonemal band. (A) Location of the BB marker SAS4-GFP (green) in relation to axoneme marker kinesin-8B-mCherry (red) in male gametocytes at 3-4 minutes and 6-8 minutes post activation, highlighting the formation of axonemes around the nucleus (Hoechst staining, blue). Scale bar = 5 μm. (B) Microgametocyte 3D models (viewed at 90^°^C rotations) from SBF-SEM data showing the axonemes forming a band around the nucleus. Axonemes are shown in yellow or blue depending on their direction of coiling around the nucleus starting at the basal body (red). Axonemes are numbered to follow each axoneme. Scale bar = 1 μm. (C) Microgametocyte 3D model from SBF-SEM data showing the nucleus (cyan) surrounded by axonemes (blue/yellow) with paired basal bodies (red). Scale bar = 500 nm. (D) Inset showing the detail of Fig 2C 3D model showing a basal body (BB - red) and nuclear pole (NP - purple) pair with their associated axonemes (blue) coiling in the same direction. Scale bar = 500 nm. (E) Bar graph showing the total length of each axoneme per microgametocyte cell (N=120 axonemes). Axoneme lengths are divided in regions – where the central pair is present (9+2, yellow) and where it is absent (9+0, green) – see text. (F) SBF-SEM data slices highlighting the presence of the outer doublets and central pair (z = 0 nm) and the presence of the outer doublets only in a neighbouring section of the same axoneme (z = 100 nm). Scale bar = 1 μm.

### Outer doublet microtubule assembly precedes central pair formation in elongating axonemes

This unique SBF-SEM 3D dataset had sufficient resolution to allow us to analyse and quantify axoneme assembly in each microgametocyte. The length of each axoneme in a cell was measured and was found to be highly variable in all cells ranging from 3 μm to 20 μm, with an average of 13.6 +/- 0.9 μm (Fig 2E). There was sufficient resolution in these datasets to distinguish clearly the outer doublets of a 9 + 2 axoneme as an electron dense circle with the central pair appearing as a central electron density within this electron dense circle (Fig 2F; z = 0 nm). This then allowed us to analyse outer doublet and central pair assembly, as sections without a central pair are easily visible (Fig 2F; z = 100 nm) and could reveal the progress of axoneme assembly at this pre-exflagellation stage. Assembly of central pair microtubules have been shown to precede outer doublet microtubule assembly during axoneme growth in the Kinetoplastida parasite *Trypanosome brucei* by ∼50nm, but in fully assembled older axonemes central pair microtubules extend to the distal end (Höög et al., 2014). All 8 axonemes had begun assembly in each of the 15 microgametocytes (Fig 2E; n = 120 axonemes,). However, all microgametocytes contained axonemes that did not have fully elongated central pair microtubules and only 40 out of 120 axonemes had a fully elongated central pair (Fig 2E). Using these criteria we conclude that most axonemes are still in the process of assembly at this stage in male gametogenesis. This finding is an important aspect to take into consideration when assessing the timing of gametogenesis and analysing axoneme mutants.

### Unique organisation of the nucleus around the axoneme of microgametes

The nucleus was reconstructed in each cell and this revealed a highly contorted 3D structure (Fig 3A; 2 cell examples – N=15 cells). There were multiple projections with BB/NPs often positioned at the end of these projections (Fig 3B; arrow). Measurements of nucleus volume revealed an average volume of 27 +/- 3.4 μm^3^. There was some variability between nuclei volume, but this was not significant. Our datasets also included cells further along the process of gametogenesis, where individual axonemes were found to be in the process of ex-flagellating (Fig 3C-G, Movie 3). Not all microgametocytes at this stage contained 8 flagella, but since all 15 microgametocytes contained 8 internally coiled axonemes (Fig 2B), it is likely that some axonemes have already ex-flagellated in these examples. In one microgametocyte there were 2 microgametes undergoing ex-flagellation and no internal axonemes suggesting that this cell was in a late-stage of ex-flagellation with 6 microgametes having already been released (Fig 3E). Despite the late stage in ex-flagellation there was still a wide variation in axoneme length in these microgametocytes (Fig 3H; exflagellating cells), while microgamete axonemes seemed less variable in length (Fig 3H; microgametes), suggesting that axoneme extension is still undergoing in microgametocytes at the ex-flagellation stage.

**Figure 3.**
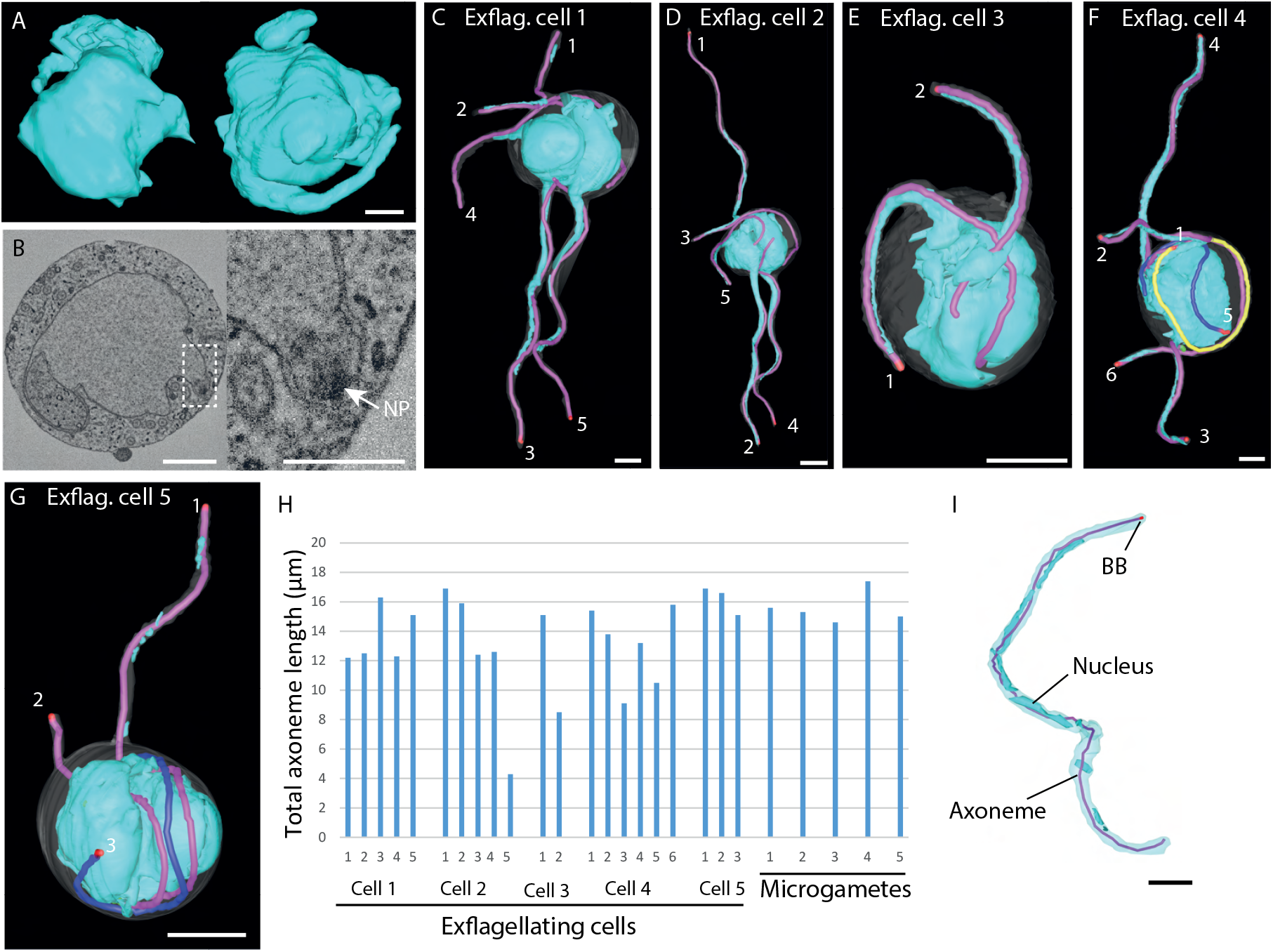
A portion of the nucleus associated with each axoneme in exflagellating microgametocytes. (A) Two different 3D models of nuclei showing the complex 3D structure and nuclear projections. Scale bar = 1 μm. (B) SBF-SEM data slice of a nucleus and the positioning in of the nuclear pole (NP) within a nuclear projection. Scale bar = 1 μm. (C-G) 3D models of microgametocytes undergoing exflagellating at different stages. Nucleus (blue), plasma membrane (white), basal body (red), exflagellating axoneme (pink), internally coiled axoneme (yellow/blue). Scale bar = 1 μm. (H) Bar chart showing the axoneme lengths of exflagellating cells (see Figure 3C-G) and microgametes. (I) Segmented free microgamete highlighting the ultrastructural features (basal body – red, axoneme – pink, plasma membrane – pale blue, nucleus – blue) with the nucleus coiling around the axoneme. Scale bar = 1 μm.

During ex-flagellation a portion of the microgametocyte nucleus containing a haploid genome is taken to form the microgamete nucleus (Rashpa & Brochet, 2022; R. Sinden et al., 1978; Zeeshan et al., 2019, 2022). However, little is known about how this genome inheritance is organised at the ultrastructural level. This was investigated further in microgametocytes undergoing ex-flagellation and in ex-flagellated microgametes. Intriguingly, our SBF-SEM imaging revealed that the nucleus is coiled around the axoneme of ex-flagellating and ex-flagellated microgametes, which has not been previously reported (Fig 3C-G, 3I).

For improved resolution of the elongated nuclear structures, we performed dual axis cellular electron tomography of portions of ex-flagellated microgametes in order to understand how this unique coiling is organised (Fig 4A, Movie 4). This clearly showed nuclear structures tightly coiled around the outer doublet microtubules of the axoneme enclosed within the flagellar membrane. There were distinctive focal electron densities within the membrane of the nucleus that was asymmetrically located close to the point where the nuclear envelope was closely associated with the outer doublet microtubules of the axoneme. This could represent the condensed chromatin indirectly ‘cross-linked’ to axonemal microtubules through the nuclear envelope (Fig 4A; asterisks, Movie 4). Each central point of each density was selected in each slice of the tomogram, which showed these densities to coil around the axoneme (Fig 4B-C). An important question is when coiling of the nucleus around the axoneme occurs. In SBF-SEM reconstructions of microgametocytes at later stages of ex-flagellation there were many examples of flagella that were partially ex-flagellated (Fig 3C-G). In the portion of flagellum still contained within the microgametocyte, we frequently observed the nucleus folding around the flagellum (Fig 4D-F). Extensive analysis using thin section transmission electron microscopy combined with cellular electron tomography revealed the nucleus interacting with the axoneme in a similar coiled arrangement (Fig 4G-H, Movie 5) as seen in the microgamete (Fig 4A). In addition, the same electron densities were seen on the inner nuclear envelope in close association with the axoneme (Fig 4G-H; asterisks). This confirms that coiling of the nuclear material starts when axonemes are still within the microgametocyte. Microgametes are motile flagella and, like all flagella, have distinct wave forms patterns. Coiling of the nuclear material around the axoneme might be important in maintaining this wave form. Therefore, having the nucleus elongated along the axoneme would be advantageous for motility. Taken together, these results suggest there is a highly organised periodic coiling of nuclear material around the axonemes within the microgamete and represents an extra-axonemal structure within microgametes of *Plasmodium berghei* which lacks IFT.

**Figure 4.**
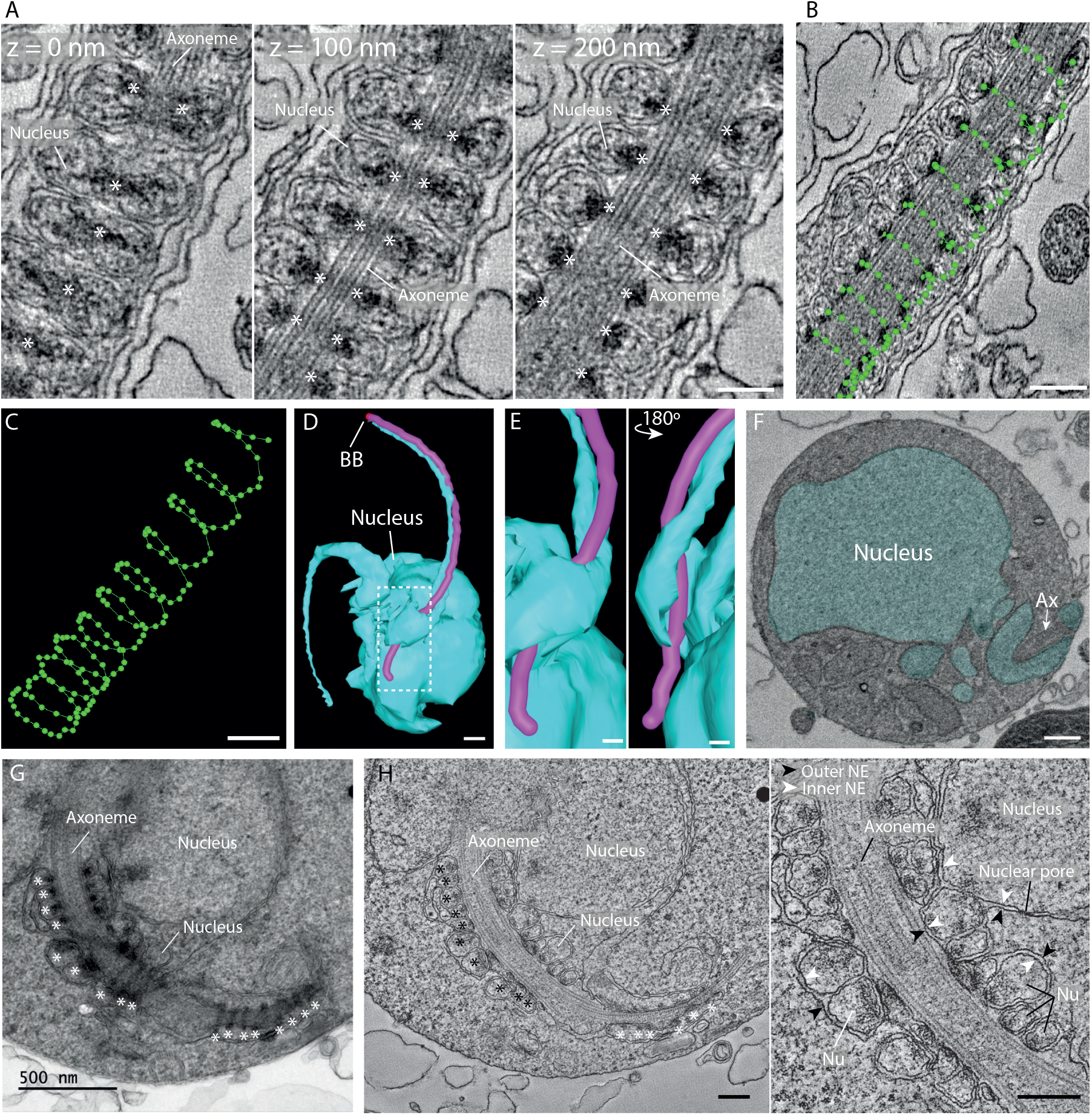
(A) Serial section tomography micrographs of a free microgamete showing the nucleus and electron dense material (asterisks) coiling around the axoneme. Scale bar = 200 nm. (B-C) 3D models of a free microgamete (shown in Fig 4A) showing the coiling of the electron dense material (green) around the axoneme. Scale bar = 200 nm. (D) 3D model of an exflagellating microgametocyte showing the nucleus (blue) folding around the axoneme (pink).Scale bar = 500 nm. (E) Higher magnification 3D model of Fig 4D showing the nucleus (blue) folding around the axoneme (pink) at 180^°^C rotations. Scale bar = 200 nm. (F) SBF-SEM data slice of the nucleus (blue) and axoneme (Ax) (arrow) shown in Figure 4D. Scale bar = 200 nm. (G) Transmission electron micrograph of a semi-thin (150 nm) section highlighting the coiling of the nucleus around the axoneme within a microgametocyte. Scale bar = 500 nm. (H) Serial section tomography micrographs showing that the nucleus is still connected to the nuclear envelope whilst coiling around the axoneme within a microgametocyte White arrows indicate the inner nuclear envelope (NE) and black arrows indicate the outer nuclear envelope (NE). Nu, nucleus. Scale bar = 200 nm.

## Conclusions

The technology and development of SBF-SEM and cellular electron tomography has enhanced our understanding of how BB-NPs are organised in *Plasmodium* during male gamete formation and revealed an axonemal band around the nucleus with axonemes coiled in different directions showing chirality within this tight band. How axoneme assembly is restricted to this central band is not understood but might be influenced by the 3D organisation between BB and NPs in directing this process, along with inherent constraints in axoneme extension. In addition, tight positioning of axonemes in a central axonemal band could be due to localised accumulation of axoneme assembly proteins within this band. Furthermore, the organisation of axonemes in opposing orientations could generate momentum and rotational power to drive ex-flagellation.

The intricate coiling of the nucleus around the axoneme described here shows a level of ultrastructural complexity within the microgamete that has not previously been reported and further work will be required to dissect the precise mechanism that drives coiling of the nucleus around the axoneme.The rapid process of microgamete generation is widely reported to be error-prone, and the detailed analysis shown here provides insights into the causes and complexity in the association between BBs/axonemes and nuclear structures in Plasmodium microgametocytes.

## Acknowledgements

We would like to thank The Oxford Brookes Centre for Bioimaging for assistance in carrying out this project. Molly Hair was funded by a Nigel Groome studentship. We thank Declan Brady at University of Nottingham for technical assistance in mice and parasite work. This work was supported by MRC UK (MR/K011782/1) to Rita Tewari and BBSRC (BB/N017609/1) and ERC advance grant funded by UKRI Frontier Science (EP/X024776/1) to Rita Tewari and Mohammed Zeeshan.

## Methodology

### Ethics statement

The animal work done at University of Nottingham passed an ethical review process and was approved by the United Kingdom Home Office. Work was carried out under UK Home Office Project Licenses (30/3248 and PDD2D5182) in accordance with the UK ‘Animals (Scientific Procedures) Act 1986’. Six- to eight-week-old female CD1 outbred mice from Charles River laboratories were used for all experiments.

### Parasite culture and gametocyte purification

*Plasmodium berghei* transgenic lines expressing SAS4-GFP (Zeeshan et al, 2022; PMID: 35550346), kinesin-8B-mCherry (Zeeshan et al, 2019; PMID: 31409625) and NDC80-mCherry (Zeeshan et al, 2020; PMID: 32501284) were injected into phenylhydrazine treated mice (Beetsma et al., 1998, PMID: 9501851). Gametocyte enrichment was achieved by sulfadiazine treatment after 2 days of infection. The blood was collected on day 4 after infection and gametocyte-infected cells were purified on a 48% v/v NycoDenz (in PBS) gradient (NycoDenz stock solution: 27.6% w/v NycoDenz in 5 mM Tris-HCl, pH 7.20, 3 mM KCl, 0.3 mM EDTA). The gametocytes were harvested from the interface and activated in ookinete medium containing xanthurenic acid.

### Generation of dual tagged parasite lines and live cell imaging

Parasite cells from lines expressing tagged SAS4-GFP (green) were mixed with cells from mCherry (red) tagged lines of kinetochore marker NDC80 (Zeeshan et al, 2020; PMID: 32501284) and axoneme marker kinesin-8B (Zeeshan et al, 2019; PMID: 31409625) in equal numbers and injected into mice. Mosquitoes were fed on these mice 4 to 5 days after infection when gametocytemia was high and were checked for oocyst development and sporozoite formation at days 14 and 21 after feeding. Infected mosquitoes were then allowed to feed on naïve mice and after 4 to 5 days the mice were examined for blood stage parasitaemia by light microscopy of Giemsa-stained blood smears. Fluorescence microscopy showed that some parasites expressed both SAS4-GFP and NDC80-mCherry; and SAS4-GFP and kinesin-8B-cherry in the resultant gametocytes. These gametocytes were purified as described above, activated and fluorescence microscopy images were captured at desired time points using a 63x oil immersion objective on a Zeiss Axio Imager M2 microscope fitted with an AxioCam ICc1 digital camera.

### Serial block face scanning electron microscopy (SBF-SEM) of culture derived gametocytes

*Plasmodium berghei* cells were fixed in 2.5% glutaraldehyde in 0.1 M phosphate buffer 15 mins after activation to allow for an asynchronous population of the different stages of gametogenesis and exflagellation to be identified and analysed. Samples were then spun and washed three times in 0.1M phosphate buffer, and post-fixed in 1% osmium tetroxide in 1.5% potassium ferrocyanide in 0.1M phosphate buffer (for 45 min, at RT, in the dark). After the first osmium step, samples were washed three times in 0.1M phosphate buffer, incubated in 1% tannic acid in 0.1M phosphate buffer for 30 mins, and then subjected to a second osmium step (2% OsO_4_ in ddH_2_O, for 30 min, at RT, and in the dark). Samples were then incubated in 2% uranyl acetate in ddH_2_O for 2h, dehydrated in acetone and embedded in TAAB 812 Hard resin (TAAB, catalogue number T030). The tips of resin blocks containing samples were trimmed and mounted onto aluminum pins using conductive epoxy glue and silver dag, and then sputter coated with a layer (10-13 nm) of gold in an Agar Auto Sputter Coater (Agar Scientific). Before SBF-SEM imaging, ultrathin sections (70 nm) of the block face were examined in a Jeol JEM 1400 Flash transmission electron microscope (JEOL), to verify sample quality. Samples were then imaged in a Merlin VP compact high resolution scanning electron microscope (Zeiss) equipped with a 3View stage (Gatan-Ametek), an OnPoint back-scattered electron detector (Gatan-Ametek), in variable pressure. The following imaging conditions were used: 3kV, 30 μm aperture, 30 pascal variable pressure, 3 nm pixel size, 4 μs pixel time, 100 nm section thickness.

### SBF-SEM segmentation and data analysis

Data were processed using the IMOD software package (Kremer et al., 1996). Briefly, image stacks were assembled, corrected (for z scaling and orientation) and aligned using eTOMO, and 3D models were produced using 3dmod. Whole individual microgametocyte cells with internally coiled axonemes (n = 15), microgametocytes undergoing exflagellation (n = 5), and free microgametes (n = 5) were identified and manually segmented from trimmed regions of original datasets. In each cell, the cell membrane, the nucleus, the axonemes with subtending basal bodies and the nuclear poles were identified based on distinctive ultrastructural features, and then segmented manually as individual objects. For the nucleus, the outer membrane was used to define the edges of segmentation, and the nuclear poles were segmented as the area that included all electron dense structures known to form the NP. Volume, surface area and length measurements for different structures were obtained in 3dmod, based on each object’s surface rendering.

### Dual axis cellular electron tomography

*Plasmodium berghei* cells were fixed in 4% glutaraldehyde in 0.1 M phosphate buffer and processed for electron microscopy.Samples were post-fixed in osmium tetroxide, treated en bloc with uranyl acetate, dehydrated and embedded in Spurr’s epoxy resin. Thin sections were stained with uranyl acetate and lead citrate prior to examination in a JEOL JEM-1400 electron microscope (JEOL, UK). Serial-section cellular electron tomography (ssET) was performed on 150 nm sections collected onto formvar-coated slot grids. Grids were mounted in a Fishione dual-axis tomography holder (Fischione instruments) and dual-axis tilt-series (55° to -55°, with 1° tilt between images) of the same area in consecutive sections were acquired using SerialEM (Mastronarde, 2005). Tomogram generation and serial tomogram joining were perfomed in ETomo (IMOD software package: https://bio3d.colorado.edu/imod/).

## Figure Legends

**Movie 1**. Serial block face scanning electron microscopy dataset. The dataset section thickness is 100 nm Z resolution. Scale bar = 2 μm.

**Movie 2**. SBF-SEM imaging and segmentation of a single microgametocyte post activation showing 8 axonemes coiling around the nucleus in two orientations. Organelles are modelled on the following colours: plasma membrane (white), nucleus (blue), basal body (red), axonemes (yellow/blue – two colours to highlight directions of coiling). Scale bar = 1 μm.

**Movie 3**. SBF-SEM imaging and 3D model of a microgamete cell undergoing exflagellation. Structures are modelled on the following colours: plasma membrane (white), nucleus (blue), basal body (red), flagella (pink). Scale bar = 1 μm.

**Movie 4**. Serial section electron tomography of a microgamete highlighting the coiling of the nucleus around the axoneme. Scale bar = 200 nm.

**Movie 5**. Serial section electron tomography of a microgametocyte highlighting the coiling of the nucleus around the axoneme. Scale bar = 200 nm.

